# Nucleosome dynamics render heterochromatin accessible in living human cells

**DOI:** 10.1101/2024.12.10.627825

**Authors:** Hemant K. Prajapati, Zhuwei Xu, Peter R. Eriksson, David J. Clark

## Abstract

The eukaryotic genome is packaged into chromatin, which is composed of a nucleosomal filament that coils up to form more compact structures. Chromatin exists in two main forms: euchromatin, which is relatively decondensed and enriched in transcriptionally active genes, and heterochromatin, which is condensed and transcriptionally repressed ^1–10^. It is widely accepted that chromatin architecture modulates DNA accessibility, restricting the access of sequence-specific, gene-regulatory, transcription factors to the genome. Here, we measure genome accessibility at all GATC sites in living human MCF7 and MCF10A cells, using an adenovirus vector to express the sequence-specific *dam* DNA adenine methyltransferase. We find that the human genome is globally accessible in living cells, unlike in isolated nuclei. Active promoters are methylated somewhat faster than gene bodies and inactive promoters. Remarkably, both constitutive and facultative heterochromatic sites are methylated only marginally more slowly than euchromatic sites. In contrast, sites in centromeric chromatin are methylated slowly and are partly inaccessible. We conclude that *all* nucleosomes in euchromatin and heterochromatin are highly dynamic in living cells, whereas nucleosomes in centromeric α-satellite chromatin are static. A dynamic architecture implies that simple occlusion of transcription factor binding sites by chromatin is unlikely to be critical for gene regulation.

---

Chromatin consists of repeating units called nucleosomes, which contain ∼147 bp of DNA coiled around a central histone octamer core. The octamer is composed of two molecules each of H2A, H2B, H3 and H4 ^11^. Nucleosomes are regularly spaced on the DNA, resembling beads on a string. This nucleosomal filament undergoes additional compaction to form euchromatin or heterochromatin. In mammalian cells, heterochromatin may be constitutive, involving the same genomic regions in all cells (typically gene-poor regions composed of short repeated sequences), such as pericentromeric and telomeric regions, or facultative, involving cell type-specific fully repressed genes ^1–10^. Heterochromatin is associated with specific post-translational histone modifications: constitutive heterochromatin is marked by H3K9me2/3, whereas facultative heterochromatin is marked by H3K27me3, although there is some overlap ^12^. Conversely, other histone marks, such as H3K4me1, H3K4me3, H3K36me3 and H3K27ac, are generally associated with euchromatin.

The highly condensed nature of heterochromatin suggests that access to the DNA may be limited or even prevented. However, large proteins and dextrans can penetrate heterochromatin domains to some extent when injected into living cells, suggesting that heterochromatin may be accessible ^13^. Furthermore, heterochromatin protein 1 (HP1), which binds to H3K9me3 in constitutive heterochromatin, is mobile in living mammalian cells ^14,15^ and transcription of repeat sequences in constitutive heterochromatin occurs at low levels ^6,16^. These data indicate that constitutive heterochromatin is at least partially accessible some of the time. Liquid-liquid phase separation may also be important in constitutive heterochromatin, resulting in exclusion of specific proteins from the heterochromatin phase ^9,17–19^. These studies have led to a more nuanced view concerning the accessibility of constitutive heterochromatin.

Facultative heterochromatin contains inactive genes that are subject to Polycomb-mediated repression and are marked by H3K27me3 (reviewed by ^10^). Genome-wide MNase-seq and ATAC-seq studies on isolated nuclei from various organisms have shown that inactive genes lack nucleosome-depleted regions (NDRs) at their promoters, unlike active genes. This observation suggests that nucleosomes prevent transcription factor binding at regulatory elements, such as promoters and enhancers, resulting in repression ^20–24^. However, inactive promoters are partially accessible in mouse liver cell nuclei ^25^. Although most transcription factors cannot access their cognate binding sites when incorporated into a nucleosome ^20^, there is a class of transcription factor, the “pioneer” factors, which bind to a nucleosomal site with high affinity ^26^. Pioneer factors may be critical for initiating the process of nucleosome removal from regulatory elements by facilitating the binding of other transcription factors and recruitment of ATP-dependent chromatin remodelers to remove or displace blocking nucleosomes ^27–29^.

These observations suggest that nucleosomes play a crucial role in gene regulation by controlling access to regulatory elements. However, they are based primarily on experiments with nuclei, which may not be representative of chromatin in living cells. Indeed, we have shown recently that the budding yeast genome is globally accessible in living cells, except for the point centromeres and the silenced loci ^30^. However, budding yeast chromatin is virtually all euchromatin, and lacks heterochromatin resembling that found in higher eukaryotes, Here, we have asked whether human euchromatin is generally accessible in living cells, like that of yeast, and whether human heterochromatin is inaccessible, as might be expected. Surprisingly, we find that both euchromatin and heterochromatin are generally accessible at the nucleosomal level in living cells, and that only particular centromeric regions have limited accessibility in vivo.

## Global accessibility in live human cells

We adapted our qDA-seq method to measure genome accessibility in human cells ^25,30^. Specifically, we used *E. coli dam* methyltransferase (Dam) as a probe for the accessibility of GATC sites in chromatin. Dam methylates the ‘A’ in GATC to ‘m^6^A’. It is challenging to express Dam without leaky expression using inducible promoter systems ^31^. Therefore, we employed an adenovirus vector to transduce Dam fused to GFP and three HA tags into MCF7 cells (a human breast cancer cell line). Following transduction, cells were collected at various time points to monitor the kinetics of genome methylation (Fig. 1a) and of Dam production (Fig. 1b). Genomic DNA was purified and digested with DpnI, a restriction enzyme that cuts at GATC only if the ‘A’ is methylated on both strands. Agarose gel analysis of the extent of DpnI digestion revealed that the genome became almost fully methylated over time (Fig. 1c; compare with purified control unmethylated MCF7 DNA completely digested with MboI, which cuts at unmethylated GATC sites). Thus, Dam can access a large fraction of the human genome.

**Fig. 1.**
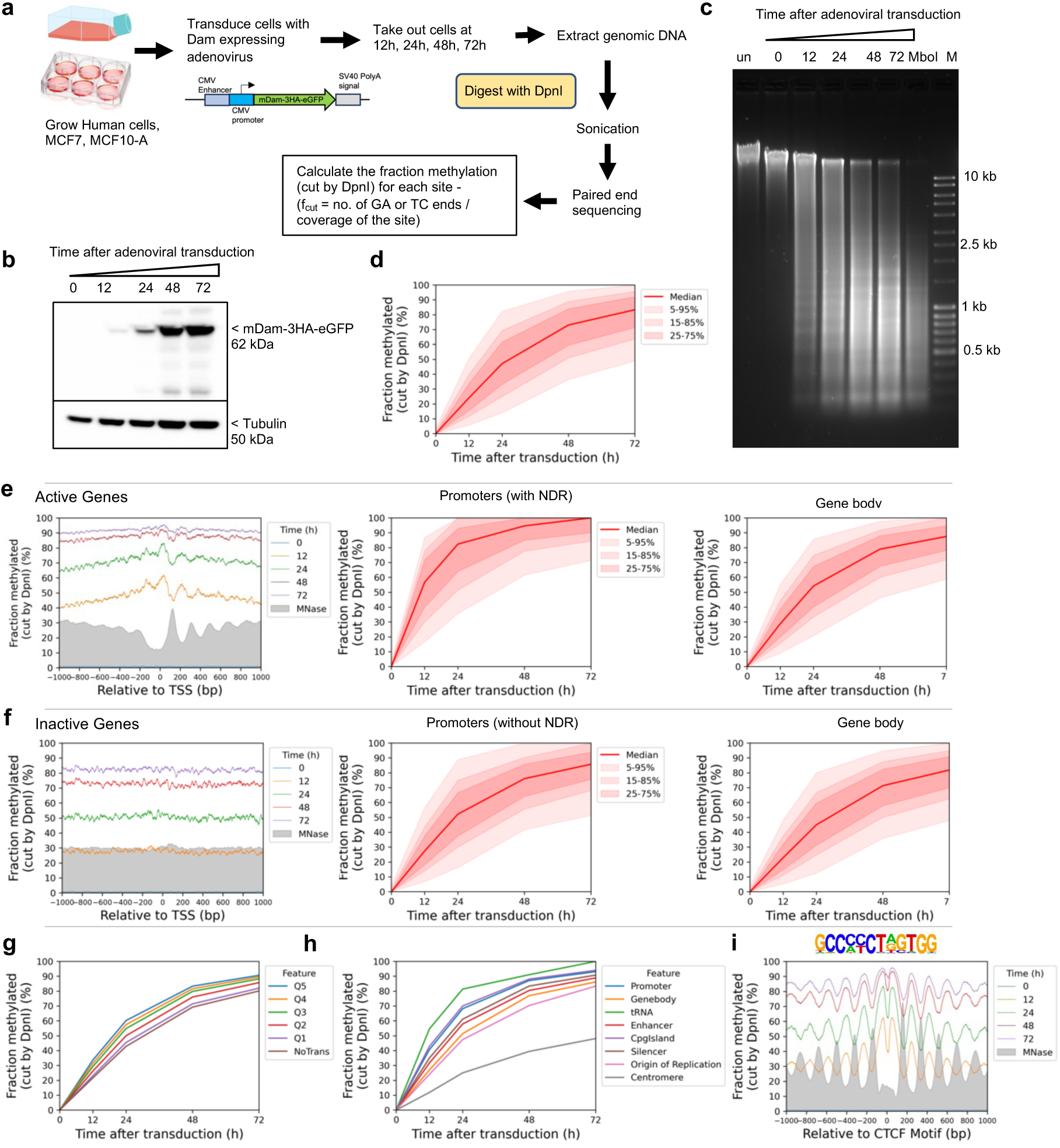
The human genome is globally accessible in living MCF7 cells. **a**, Schematic of adenovirus transduction and time course experiment to express Dam methylase in live cells. **b,** Anti-HA immunoblot to detect Dam-3HA-eGFP expression in MCF7 cells. **c,** Agarose gel electrophoresis of DpnI-digested genomic DNA purified from MCF7 cells as a function of time of adenovirus treatment. ‘un’, undigested genomic DNA; ‘MboI’, DNA from non-transduced cells digested with MboI; M, DNA size marker. **d,** Almost complete methylation of GATC sites in MCF7 cells after transduction. Red line and shading: median GATC site methylation with data range indicated. **e, f,** Nucleosome phasing with respect to the TSS for active and inactive genes, as defined by ATAC-seq ^32^. Grey profile: nucleosome dyad distribution in nuclei (MNase-seq data for MCF7 cells arbitrarily normalised to 30%). **g,** The effect of transcriptional activity on methylation rate. Active genes were divided into quintiles Q1 to Q5 based on increasing transcriptional activity (Q5 is the highest) using RNA-seq data from ^32^; methylation of the median GATC site in each quintile is shown. Inactive genes are treated as a single separate group (’NoTrans’). **h,** Median GATC methylation for various genomic regions. **i,** Nucleosome phasing around CTCF motifs using the motif shown.

The conclusion that most of the genome is accessible is further supported by genomic analysis of GATC sites (Fig. 1d). The DpnI-digested DNA was sonicated to small fragments and subjected to paired-end sequencing. For each GATC site, the fraction methylated is calculated as the number of left or right DNA fragment ends divided by the coverage of that GATC site (Fig. 1a; see Methods). We calculated the methylation kinetics for each of ∼5.9 million GATC sites in the human genome. To visualise the data for all GATC sites, we plotted the fraction methylated for the median GATC site, and for all sites within the 5%-95% methylated range, as a function of time after transduction (Fig. 1d). The median GATC site was ∼80% methylated after 72 h and still trending upwards; 90% of all GATC sites show a similar trend.

We compared the methylation kinetics of transcriptionally active and inactive genes. Analysis of published ATAC-seq data for MCF7 cells ^32^ (Extended Data Fig. 1a) delineated two gene classes: one with high ATAC signal at the promoter, indicating the presence of an NDR, and one with low or no ATAC signal, indicating the absence of an NDR (Extended Data Fig. 1a). To confirm this interpretation, we performed MNase-seq on MCF7 nuclei and sorted the genes according to ATAC signal (Extended Data Fig. 1a). We observed a clear correlation between ATAC signal and the presence of a promoter NDR. Published gene expression (RNA-seq) data for MCF7 cells ^32^ also correlate with ATAC signal (Extended Data Fig. 1a).

We examined GATC site methylation at active and inactive promoters by plotting the mean GATC site methylation as a function of distance from the transcription start site (TSS). After 12 h of transduction, active genes show a weak nucleosome phasing signal that is exactly out of phase with our nucleosome dyad data (MNase-seq) for MCF7 nuclei (grey profile) (Fig. 1e; Extended Data Fig. 1b). This suggests that Dam methylation of the linkers and in promoter NDRs is slightly faster than methylation within the first (+1) and second (+2) nucleosomes. Mean methylation increases with time, reaching ∼90% by 72 h. Promoter NDRs are methylated faster than gene bodies (Fig. 1e). Inactive genes show the same trend, but are methylated slightly more slowly, reaching ∼83% after 72 h (Fig. 1f; Extended Data Fig. 1b). The methylation rate is almost uniform across inactive promoter regions, with no phasing; promoters and gene bodies are methylated at almost the same rate, consistent with the MNase-seq data (Fig. 1f). Nucleosome positioning appears to be essentially random around inactive promoters. We conclude that both active and inactive promoter regions are almost entirely accessible to Dam in vivo.

To determine whether higher transcription renders genes more accessible, we divided the active genes into quintiles according to their mRNA levels in MCF7 cells ^32^. Quintile 5 has the most active genes; inactive genes were placed in a separate group. We observed a small but reproducible trend of increasing median methylation with increasing transcriptional activity; the most active genes were methylated marginally faster than the least active genes (Fig. 1g; Extended Data Fig. 1c). Thus, higher transcription correlates with a modest increase in methylation rate. However, the overarching conclusion is that both active and inactive genes are accessible to Dam.

## Slow, limited methylation at centromeres

To measure the accessibility of other genomic regions, we compared median methylation rates for GATC sites in promoters, gene bodies, tRNA genes, enhancers, CpG islands, silencers, replication origins and centromeres (Fig. 1h) using the hg38 genome annotations ^33^. All regions are methylated at similar rates and to high levels, similar to gene bodies, except for the tRNA genes, which are methylated even faster, and the centromeres, which are methylated much more slowly and appear to be reaching a limit (Fig. 1h; Extended Data Fig. 1d,e). We quantified the median methylation rates for the various regions relative to the median for all genomic GATC sites by plotting the log of the unmethylated fraction as a function of time after transduction (Extended Data Fig. 1f). Relative methylation rates for promoters, enhancers, silencers and CpG islands were all slightly faster (1.2 to 1.7x; see replicates) than gene bodies and replication origins (1.0 to 1.1x). tRNA genes were methylated ∼2x faster, whereas centromeres were methylated ∼3x more slowly (0.3/0.4x the genomic median). Thus, the range in median methylation rate is ∼8-fold, from centromeres (slowest) to tRNA genes (fastest). We conclude that all genomic regions examined are fully accessible, except for the centromeres.

To search for large regions of relatively inaccessible chromatin at the chromosome level, we constructed a heat map showing the mean methylation rate for each 100 kb window along each chromosome. The T2T (Telomere to Telomere) human genome was used for this analysis because the centromeric regions have been thoroughly annotated ^34^. The rate was calculated as above, using the mean methylation for all GATC sites in each window for each time point. We found that windows of slow methylation tend to cluster predominantly at a few specific regions in each chromosome (Fig. 2a). Most of these windows are situated within centromeres (Fig. 2a; red rectangles indicate centromeres). The T2T genome has revealed the complexity of human centromeres, which make up ∼6% of the genome and include many different repeats, including α-satellite repeats, human satellites (HSat1 to 5), β-satellite (βSat), γ-satellite (γSat), as well as non-satellite DNA ^35^ (Fig. 2b). The α-satellite repeats comprise variants of a ∼171 bp repeat, which can be divided into 20 supra-chromosomal families. Active α-satellite repeats are enriched in centromeric histone H3 (CENP-A) and associate with the kinetochore. All of these centromeric regions are methylated more slowly (0.4x to 0.7x) than the genomic average (1.0) (Fig. 2c, d); α-satellite and HSats exhibit the slowest methylation rates (Fig. 2d; Extended Data Fig. 2). We examined the methylation rates within the CENP-A-enriched active α-satellite SF1, SF2, SF3 and SF01 supra-chromosomal families (Fig. 2e,f). The SF1 family has the slowest methylation rate (0.2x) and reaches a limit at < 30% median methylation, indicating that a GATC site located in SF1 α-satellite DNA is inaccessible in a large fraction of cells. In summary, centromeric GATC sites are methylated more slowly than other genomic regions in living cells (Fig. 2f). This is particularly true of the active α-satellite repeats (especially the SF1 family), which are associated with CENP-A-containing nucleosomes, and are only partially accessible in vivo.

**Fig. 2.**
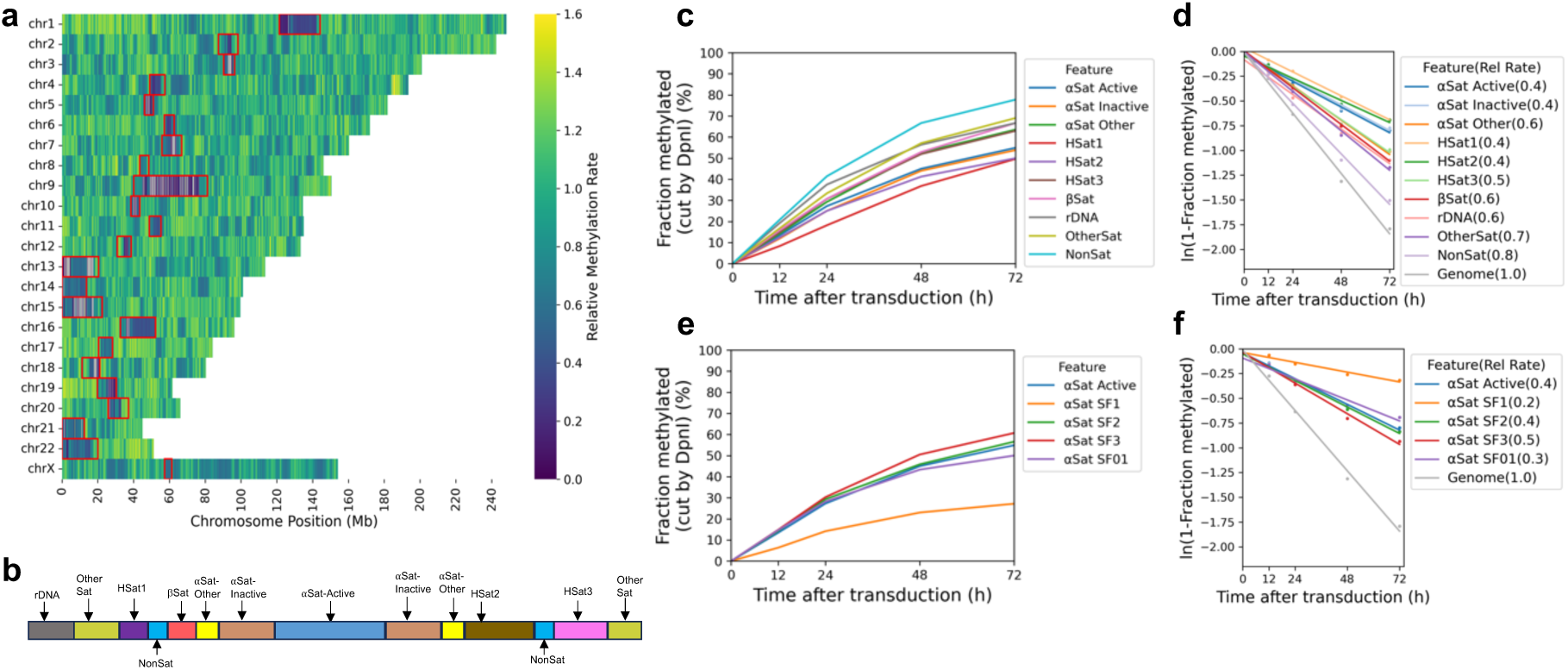
Centromeres are methylated slower and reach a limit, unlike other genomic regions. **a**, Heat map showing the variation in methylation rate at the chromosomal level. The average methylation rate was calculated for all GATC sites in each 100 kb window in the T2T genome by plotting ‘ln (1-fraction methylated)’ against time after adenovirus transduction, and then normalised to the genomic average rate to obtain relative rates. Red rectangles: centromeric regions. **b,** Schematic of the organisation of the various centromeric elements defined in the T2T genome assembly (adapted from ^35^). Divergent α-Sat and monomeric α-Sat were combined as “α-Sat other”. **c,** Median GATC methylation time courses and **d,** methylation rates for the various centromeric elements. **e,** Median GATC methylation time courses and **f,** methylation rates for the various active supra-chromosomal α-satellite families.

## TAD boundary chromatin is accessible

The loop organisation of chromatin depends on the CTCF transcription factor/insulator-binding protein and cohesin, which together define TAD boundaries ^36,37^. Consequently, CTCF binding is expected to be quite stable, which is consistent with the exceptionally good nucleosome phasing observed on both sides of CTCF binding sites (Fu et al., 2008; Wiechens et al., 2016). We examined methylation patterns at and around CTCF sites (Fig. 1i). We detected very strong phasing around CTCF motifs in our Dam data, which is out of phase with nucleosome dyads, as expected (cf. the MNase-seq dyad plot, grey profile). Although phasing is strong, the nucleosomal DNA is still accessible to Dam, since the mean methylation level increases with time, reaching ∼85% after 72 h (Fig. 1i; Extended Data Fig. 1g). The plots also imply that the CTCF site is similarly accessible to Dam, since methylation increases to high levels at the motif, even though the motif is in a small trough in the methylation profiles (Fig. 1i). However, we note that the CTCF motif itself does not contain a GATC site; it is therefore unclear whether it is protected from methylation.

## Heterochromatin is accessible in live cells

We asked whether heterochromatin, as defined by specific histone marks, is accessible in living cells using published ChIP-seq data for MCF7 cells ^38^. We grouped all GATC sites located in H3K9me3 peaks (constitutive heterochromatin) or in H3K27me3 peaks (facultative heterochromatin) and compared their methylation with GATC sites associated with euchromatin marks (H3K4me1, H3K4me3, H3K27ac and H3K36me3). We observed that GATC sites in euchromatin are methylated faster than those associated with heterochromatin (Fig. 3a; Extended Data Fig. 3a). However, the rate difference is no more than ∼2-fold: 1.2x - 1.6x the genomic average for euchromatin; 0.8x and 0.9x for the two heterochromatic states (Fig. 3a; Extended Data Fig. 3a). Most importantly, these heterochromatic regions are fully accessible to Dam.

**Fig. 3.**
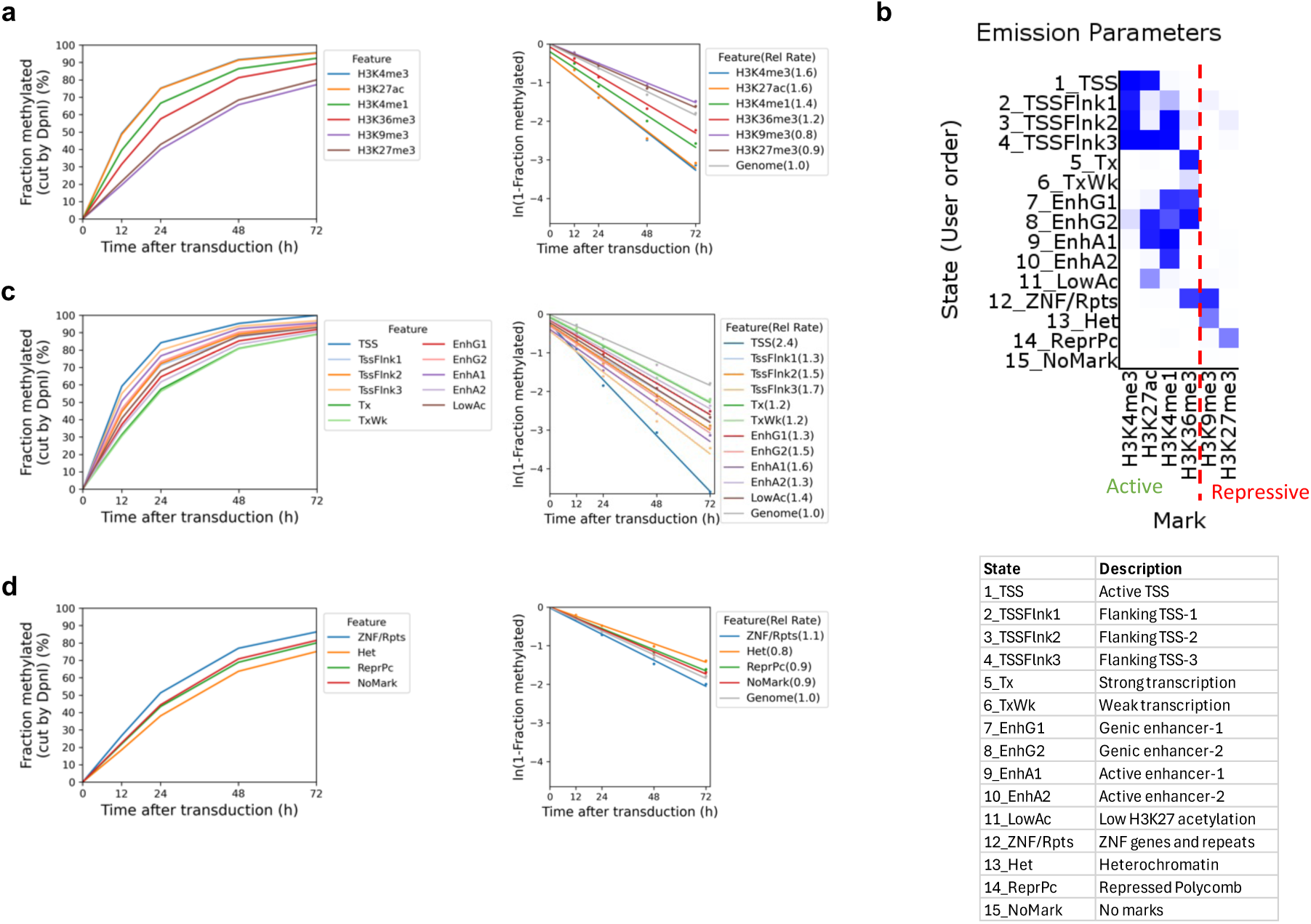
Heterochromatin is accessible but methylated at a slower rate than euchromatin. **a**, Methylation time courses and methylation rates for the median GATC site in regions with active histone marks (H3K4me1, H3K4me3, H3K27ac, H3K36me3) and inactive histone marks (H3K9me3 and H3K27me3). **b,** ChromHMM model defining 15 epigenetic states in MCF7 cells, defined as active or inactive chromatin based on the histone marks. **c,** Methylation time courses and methylation rates for the median GATC site in the active chromatin (euchromatin) states defined by the HMM model. **d,** Methylation time courses and methylation rates for the median GATC site in the inactive chromatin (heterochromatin) states defined by the HMM model.

In a more sophisticated approach, we compared methylation rates in euchromatin and heterochromatin using a 15-state epigenetic ChromHMM model ^39^, which we derived using the same MCF7 ChIP-seq data. ChromHMM models identify genomic regions associated with various combinations of histone marks. Our ChromHMM model identifies 11 euchromatin states based on the presence of H3K4me1, H3K4me3, H3K27ac and/or H3K36me3 (Fig. 3b). These are: transcription start sites (TSS; state 1), TSS-flanking regions (states 2, 3 and 4), transcriptionally active regions (state 5), weakly active regions (state 6), four types of enhancer (states 7, 8, 9 and 10), and regions with low levels of H3K27ac (state 11). Our model also defines two heterochromatic states: constitutive (H3K9me3; state 13) and polycomb-repressed (facultative) (H3K27me3; state 14). Some chromatin is in a bivalent state, characterised by the presence of both the active H3K36me3 mark and the inactive H3K9me3 mark (state 12). State 15 has none of the histone marks for which we have data and accounts for ∼56% of the MCF7 genome. We plotted the methylation kinetics for the active and repressed chromatin states separately for ease of comparison (Fig. 3c,d; Extended Data Fig. 3b). The repressed states are methylated more slowly than the active states, but importantly, even the repressed heterochromatin states trend toward complete methylation (compare Figs. 3c and 3d).

With the exception of state 1 (TSS), the actual methylation rate differences are not large, ranging from ∼1.7x the median genomic rate for most of the euchromatin states to ∼0.8x the median genomic rate for the heterochromatin states (compare Figs. 3c and 3d). The relative methylation rate for GATC sites in state 1 (H3K4me3 and H3K27ac) is relatively high (2.4x and 4.6x for biological replicates), consistent with the inclusion of NDRs in this state (Fig. 3c,d; Extended Data Fig. 3b). We conclude that GATC sites located in heterochromatin are accessible in living cells.

## Accessibility is not due to replication

We considered the possibility that global genome accessibility in living MCF7 cells might be due to DNA replication. It proved technically challenging to synchronise and maintain MCF7 cells in the G1 phase of the cell cycle prior to transduction. We were also unable to obtain fully confluent MCF7 cells. Consequently, to confirm our findings more broadly and to test the possible role of replication, we performed the same experiment using MCF10A cells, a normal human breast epithelial cell line. Time course experiments after transduction of MCF10A cells with the same Dam-expressing adenovirus produced similar results to those obtained for MCF7 cells (Extended Data Fig. 4). Thus, the genome is also globally accessible in normal MCF10A cells, suggesting that this accessibility is not due to the cancerous nature of MCF7 cells. We grew MCF10A cells to confluence, when the cells cease to replicate their DNA, and then repeated the transduction time course (Extended Data Fig. 5). We observed similarly high accessibility in these arrested cells. We conclude that DNA replication is not a major contributor to genome accessibility.

## The X-chromosome is methylated slowly

Dosage compensation results in inactivation of one of the two copies of the X-chromosome in female cells and its condensation into heterochromatin (the ‘Barr body’; reviewed by ^40^). We reasoned that the active X-chromosome would be methylated faster than the inactive X, predicting an intermediate average methylation rate relative to the autosomes, because our method cannot distinguish between the two copies. Since MCF7 cells have aberrant ploidy, whereas MCF10A cells have normal ploidy, we focused our analysis on MCF10A cells. Using the average methylation rate data for 100 kb chromosome windows (Extended Data Fig. 4i), we plotted the fraction of windows with a given relative rate for each of the 23 chromosomes (Extended Data Figs. 4l, 5k). This approach separates out the centromeric chromatin regions, which are methylated more slowly in all chromosomes. We observed that the mean relative methylation rate of the most common autosomal 100 kb window is 1.05, whereas that of the X-chromosome is 0.85. This result is consistent with slower methylation of the inactive X-chromosome due to its heterochromatic nature.

## Limited accessibility in isolated nuclei

It has been shown previously that genome accessibility is limited in nuclei isolated from both yeast and mouse liver cells ^25,30,41^. To determine whether this is also true for MCF7, we treated isolated MCF7 nuclei with increasing concentrations of purified Dam enzyme. After a 30 min incubation at 37°C, genomic DNA was purified and digested with DpnI (Fig. 4a). In comparison with the fully digested unmethylated control DNA (’MboI’ lane), nuclei samples showed incomplete DpnI digestion even at the highest Dam concentration. A clear nucleosome ladder pattern is observed in all of the Dam-treated samples, suggesting that Dam methylates linker DNA, but not nucleosomal DNA in nuclei. This result contrasts with the almost complete methylation observed in living cells (cf. Fig. 1c).

**Fig. 4.**
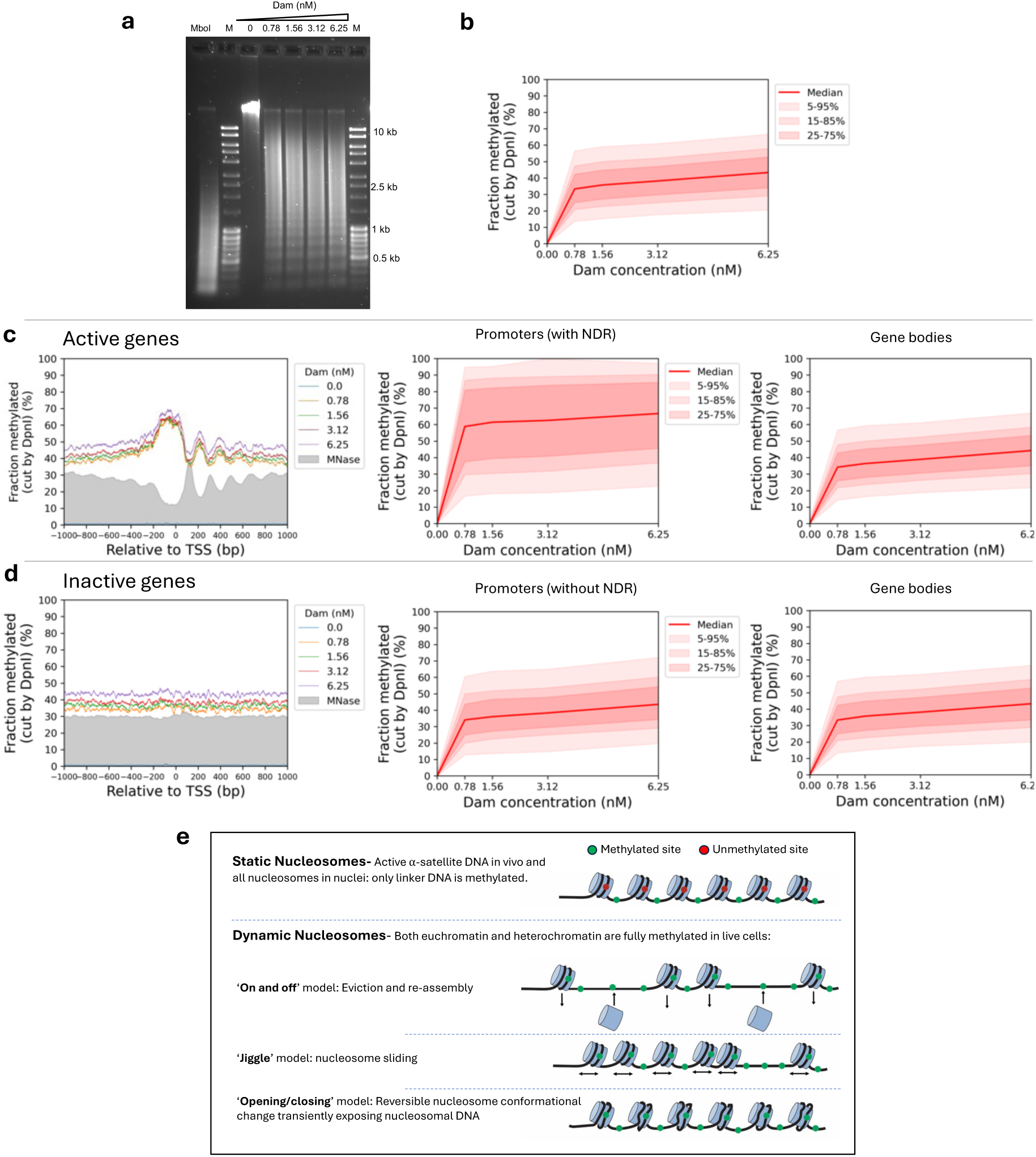
Genome accessibility is limited in isolated MCF7 nuclei. **a**, Agarose gel electrophoresis of DpnI-digested genomic DNA purified from nuclei treated with increasing amounts of Dam. M, DNA marker. MboI, unmethylated control DNA fully digested at GATC sites by MboI. **b,** Methylation of all the GATC sites in the human genome as a function of Dam concentration. Red line: methylation of the median GATC site; shading indicates the data range. **c,** Active genes: Nucleosome phasing and methylation of the median GATC site in promoters or gene bodies as a function of Dam concentration. Red line and shading: median GATC site methylation with data range indicated. Grey profile: nucleosome dyad distribution in nuclei (MNase-seq data for MCF7 cells arbitrarily normalised to 30%). **d,** The same analysis for inactive genes. **e,** Possible mechanisms for generating accessibility in living cells (based on the known activities of various ATP-dependent chromatin remodelers).

Genomic analysis confirmed that methylation is limited in nuclei (Fig. 4b; Extended Data Fig. 6). The median of all genomic GATC sites reached a plateau at ∼38% methylation, indicating that the median GATC site is accessible in ∼38% of nuclei and inaccessible in the remaining ∼62% of nuclei. Examination of Dam methylation around the TSS revealed that active genes display improved nucleosome phasing relative to living cells (compare Fig. 4c with Fig. 1e). Methylation of active genes in nuclei reaches a limit at ∼45% in the regions flanking the promoter NDR and a limit of ∼67% in the promoter NDR (Fig. 4c). In contrast, methylation of inactive genes in nuclei is uniformly limited to ∼45% over the entire region, including the inactive promoters, which are not nucleosome-depleted (Fig. 4d; MNase-seq data: grey profile). Methylation is also limited around CTCF motifs in nuclei (Extended Data Fig. 6d). The NDR associated with CTCF motifs is ∼70% accessible in nuclei and flanked by well-phased nucleosomes. Euchromatin and heterochromatin regions have very similar accessibilities in nuclei (Extended Data Fig. 3c). Application of our ChromHMM model to the nuclei data revealed that the active states reach a limit methylation of 40% to 50%, except for the TSS state which reaches ∼60% (Extended Data Fig. 3d). The TSS state is higher because it includes promoter NDRs. All of the inactive states show almost identical limit median methylation, at 35%-45%, which is slightly lower than the median limit for the active states. We conclude that accessibility is severely limited in nuclei, unlike in living cells.

Analysis of methylation at the various types of centromeric repeat in nuclei indicated that the limit median methylation ranged from ∼45% for non-satellite and other satellite repeats, similar to that observed for gene bodies in nuclei, down to ∼25%-35% for HSat1 and active and inactive α-satellites (Extended Data Fig. 7a,b). The active α-satellite SF2, SF3 and SF01 families were methylated to 25%-30% maximum (Extended Data Fig. 7c,d). Furthermore, the α-satellite SF1 family reached a limit methylation at only ∼15%, indicating that GATC sites in SF1 repeats are mostly inaccessible in nuclei. The methylation kinetics of these centromeric regions in nuclei are similar to those observed in living cells, which also tend toward a limit (Fig. 2c, e), although the methylation levels reached at α-satellite repeats in living cells are generally higher than in nuclei. Notably, the SF1 α-satellite family is the slowest methylating region observed in vivo (Fig. 2f) and the least accessible in isolated nuclei (Extended Data Fig. 7c,d).

## Human chromatin is dynamic in live cells

We have measured the accessibility of GATC sites genome-wide in vivo. We expected to find that human euchromatin would resemble yeast chromatin in being globally accessible and this is indeed the case. We proposed that yeast nucleosomes are in continuous flux in living cells, but not in nuclei, where they are static ^30^. Such a flux may occur either through nucleosome removal and replacement, or by sliding along the DNA, and/or through reversible conformational changes (Fig. 4e). It is likely that all three mechanisms occur through the agencies of multiple ATP-dependent chromatin remodelers. This flux renders the underlying DNA open to methylation by Dam and, by inference, to sequence-specific transcription factors.

The general accessibility of both active and inactive genes to Dam suggests that the widely accepted model that inactive genes are inactive because transcription factor binding sites in their promoters are blocked by nucleosomes may no longer be tenable. It seems unlikely that nucleosomes present a permanent block to transcription factor binding, thus maintaining genes in the repressed state. The role of pioneer factors, which have similar affinity for nucleosomal and non-nucleosomal sites, is unclear, given nucleosome flux. However, pioneer factors would be predicted to bind faster, since they do not have to wait for nucleosome dynamics to expose their cognate sites. Pioneer factors might also be important for initiating nucleosome dynamics. Alternatively, gene regulation may occur primarily through regulation of transcription factor gene expression, location (e.g. retention in the cytosol) or activity (e.g. post-translational modifications and allosteric effects), and through non-coding RNA expression.

We also expected that human heterochromatin might either be resistant to Dam methylation, because of its highly condensed state, or that it might be similar to chromatin in nuclei, with immobile nucleosomes, such that only linkers are methylated. Instead, we observed a trend towards complete methylation, albeit at a somewhat slower rate than for euchromatin. The relatively slow methylation of heterochromatin may be due to a combination of various factors, perhaps including slower nucleosome flux relative to euchromatin, the highly condensed nature of heterochromatin, and the presence of heterochromatin proteins (e.g., HP1 or Polycomb complexes). Nevertheless, both constitutive and facultative heterochromatin are generally accessible in living human cells. This observation is inconsistent with models proposing that heterochromatin condensation prevents access to the DNA it contains, resulting in gene repression (discussed by ^7^). Our data show that heterochromatic DNA is generally accessible and highly dynamic at the nucleosomal level in live cells, unlike in isolated nuclei.

The only genomic regions displaying limited accessibility in living human cells are the centromeric active α-satellite repeats. These elements are methylated slowly relative to other genomic regions and, unlike those regions, reach a limit at 50%-60% methylation. The SF1 α-satellite repeats are methylated even more slowly, reaching a limit at only ∼30% methylation. The nucleosomes in active α-satellite repeats are enriched in centromeric H3 (CENP-A) and so resemble yeast centromeric nucleosomes in their resistance to methylation in vivo ^30^. The limited accessibility of centromeric active α-satellite repeats in vivo is similar to that observed for all genomic regions in isolated nuclei. This observation suggests that centromeric chromatin in live cells is static, not dynamic, with little or no nucleosome flux, such that linkers are methylated and nucleosomal DNA is protected (Fig. 4e).

In summary, we have measured the accessibility of the human genome in living cells. We find that the genome is generally accessible at the nucleosomal level, including classical heterochromatin regions marked by H3K9me3 or H3K27me3. The exception is the centromeric active α-satellite repeats, which exhibit limited accessibility. We propose that nucleosome flux creates a genome-wide open chromatin environment, in which the DNA is packaged but still accessible, facilitating the search for cognate sites by sequence-specific transcription factors.

## Methods

### Cell culture

MCF7 cells (ATCC HTB-22) were cultured in RPMI 1640 with L-glutamine (Corning 10-040-CV) supplemented with 10% Fetal Bovine Serum (FBS) (Corning 35-010-CV) and 1% penicillin/ streptomycin (Gibco 15140-148) at 37°C with 5% CO_2_/95% air in a humidified incubator. MCF10A cells (ATCC CRL-10317) were cultured in DMEM/F12 (Gibco 11320-033) supplemented with 5% horse serum (Gibco 26050070), 10 µg/ml insulin (Gibco 12585-014), 20 ng/ml epidermal growth factor (Gibco PHG0311), 500 ng/ml hydrocortisone (Sigma H0888), 100 ng/ml cholera toxin (Sigma C80520), 1% penicillin/streptomycin and 2 mM L-glutamine (Gibco 25030149). Confluent MCF10A cells were obtained by culturing in a 12-well plate for 46 h in the same medium and then transduced in medium lacking all growth factors except horse serum.

### Adenovirus transduction and Dam methylation in living cells

The Dam-3HA-eGFP cassette, codon-optimised for mouse, was constructed by gene synthesis (Thermo Fisher GeneArt) (sequence available on request). The Dam expression cassette was expressed from a CMV promoter in an adenovirus vector (human adenovirus type 5 (dE1/E3); Vector Biolabs), packaged, amplified and purified by Vector Biolabs. The final viral yield was 4.2 x 10^10^ plaque-forming units (pfu) per ml, equivalent to approximately 10^12^ viral particles per ml. About a million cells were seeded in each well of a 6-well plate one day prior to adenovirus transduction. The next day, the cells were counted and the amount of adenovirus required to achieve a multiplicity of infection (MOI) of 1000 was pre-incubated at 37°C for 30 min to improve transfer efficiency ^42^. The virus was mixed with 300 µl per well of serum-free, antibiotic-free medium (RPMI 1640 for MCF7; DMEM for MCF10A) and incubated for 5-10 min at room temperature. Meanwhile, the culture medium was aspirated from the 6-well plate and 450 µl of complete medium containing FBS was added. The transduction mixture was added to each well, mixed by gently swirling the plate a few times, and incubated at 37°C for 4 h. Then 1.8 ml of complete culture medium containing serum was added to each well.

Confluent MCF10A cells were treated slightly differently: a 12-well plate and an MOI of 2000 was used. The virus was mixed with 136 µl DMEM per well and incubated for 5-10 min at room temperature. Meanwhile, the culture medium was aspirated and 204 µl of complete medium containing FBS was added. The transduction mixture was added to each well, mixed and incubated at 37°C for 8 h. Then 1 ml of complete culture medium containing serum was added to each well.

The time course started at this point, with cells harvested after 12 h, 24 h, 48 h and 72 h (for MCF10A the time points were 12 h, 24 h, 36 h and 48 h). Cells were detached from the well by washing with 1 ml PBS and then incubating in 0.4 ml 0.25% (w/v) trypsin-0.53 mM EDTA solution (ATCC 30-2101) at 37°C for 5 min (15 min for MCF10A). Next, 1 ml medium was added to the well, cells were collected by centrifugation at 100 g for 5 min, after which the medium was aspirated, and the cells were resuspended in 1 ml medium. The cells were counted, divided up for DNA and protein extraction, quickly frozen on dry ice, and stored at -80°C. DNA extraction was performed using the PureLink Genomic DNA Mini Kit (Invitrogen) according to the manufacturer’s guidelines. Purified genomic DNA (1.2 - 1.5 µg) was digested with 10 units of DpnI (New England Biolabs (NEB) R0176L) in NEB CutSmart buffer for 2 h at 37°C.

### Immunoblotting

A pellet containing 0.3-0.5 million cells was resuspended in 0.25 to 0.4 ml of 1x lithium dodecyl sulfate buffer (Invitrogen NP0007) supplemented with 0.2 M 2-mercaptoethanol and heated for 5 min at 99°C; 10 µl was loaded on to each of two 4-12% bis-Tris polyacrylamide gels (Invitrogen NP0336) and run using MOPS/SDS running buffer (Invitrogen). Transfer of proteins to a membrane and signal development with horseradish peroxidase-conjugated anti-HA (3F10; Roche 12013819001) or anti-tubulin (Abcam ab-185067) antibodies were performed as described ^30^.

### FACS analysis

Propidium iodide staining and flow cytometric DNA analysis of MCF7 and MCF10A cells were performed as described (Mullen (2004)). Cells (0.1-0.2 million) were resuspended in 50 µl cold buffer (250 mM sucrose, 40 mM trisodium citrate, 5% v/v DMSO) and frozen at -80°C. For FACS, cells were thawed and 200 µl of ice-cold Solution A (0.03 mg/ml trypsin, 3.4 mM trisodium citrate, 0.1% v/v NP-40, 1.5 mM spermine tetrahydrochloride, 0.5 mM Tris-HCl pH 7.6) was added. The mixture was incubated at room temperature for 5 min. Subsequently, 100 µl of ice-cold Solution B (0.5 mg/ml trypsin inhibitor, 0.1 mg/ml RNase A, 3.4 mM trisodium citrate, 0.1% v/v NP-40, 1.5 mM spermine tetrahydrochloride, 0.5 mM Tris-HCl pH 7.6) was added and incubated for another 5 min at room temperature. Finally, 20 µl propidium iodide at 1 mg/ml (Invitrogen P3566) was added and incubated at room temperature in the dark to prevent photobleaching. The cells were analysed using a FACSCalibur flow cytometer (Becton Dickinson) and Cell Quest Pro software, following the manufacturer’s instructions.

### Dam methylation of isolated nuclei

MCF7 cells were cultured in complete medium in a 75 cm² flask and re-passaged into a new flask after 2-3 days of growth. When the cells reached approximately 80% confluency, they were trypsinized and harvested. To extract nuclei, a pellet of 3 - 4 million cells was resuspended in 2 ml Buffer A (15 mM Tris-HCl pH 8.0, 15 mM NaCl, 60 mM KCl, 1.5 mM EDTA, 0.5 mM spermidine, 15 mM β-mercaptoethanol, and protease inhibitors) with 0.03% NP-40. The mixture was gently but thoroughly mixed by pipetting and incubated on ice for 10 min, inverting the tube 2 or 3 times during the incubation. The lysate was centrifuged at 500 g for 2 min at 4°C, and the supernatant was removed. The nuclei were washed with 1 ml Buffer A. The nuclei were resuspended in 1 ml Buffer A supplemented with fresh S-adenosylmethionine to 0.5 mM, and divided into five 200 µl aliquots. Dam methyltransferase (NEB M0222B-HC2 at 40 U/µl; 8 µg Dam/ml) was added to the aliquots of nuclei: 0, 25, 50, 100, and 200 units (0, 0.8, 1.6, 3.1, 6.3 nM, respectively), gently mixed, and incubated for 30 min at 37°C. Genomic DNA was extracted using the PureLink Genomic DNA Kit (Invitrogen 2666617). Finally, 1.2 to 1.5 µg purified genomic DNA was digested with 10 units of DpnI as above.

### MNase-seq

MNase (Worthington LS004798) was dissolved to 10 units/µl in 5 mM Na-phosphate buffer pH 7.0, 0.025 mM CaCl2, aliquotted out, and stored at -80°C. MCF7 cells (3 to 4 million) were resuspended in 2 ml Buffer B (15 mM Tris-HCl, pH 8.0, 15 mM NaCl, 60 mM KCl, 1 mM EDTA, 2 mM CaCl_2_, 0.5 mM spermidine, 0.03% NP-40, 15 mM 2-mercaptoethanol and protease inhibitors). The cells were gently lysed by pipetting and incubated on ice for 10 min, inverting the tube 2-3 times during incubation. The lysate was centrifuged at 500 g for 2 min at 4°C and the supernatant was removed. The nuclei were washed with 1 ml Buffer B without NP40 and resuspended in 1.3 ml Buffer B without NP40. MNase was added to six tubes of 200 µl nuclei, as follows: 12.5 U, 25 U, 50 U, 100 U, 200 U and 400 U, gently mixed, and incubated for 3 min at 25°C. MNase-digested DNA was purified using the PureLink Genomic DNA kit (Invitrogen 2666617) and analysed in an agarose gel. For accurate and even nucleosome mapping, we chose digests with a dominant band at ∼150 bp corresponding to >80% of the DNA (typically 25 U, 50 U and 100 U), prepared paired-end libraries, and performed low-coverage sequencing to identify the digest with the most optimal DNA fragment length distribution ^43^. The 25 U sample (Replicate 1) and the 50 U sample (Replicate 2) were chosen for high-coverage sequencing.

### Library preparation for paired-end Illumina sequencing

For both nuclei and live cell experiments, DpnI-digested genomic DNA was purified using 1.8 vol. AMPure XP beads (Beckman). Paired-end libraries were prepared as described ^30^ except for the sonication step, in which the DNA was fragmented using a Covaris ME220 ultrasonicator (350 bp program, peak power, 50 W; duty factor 10%; 1,000 cycles per burst; average power 5; total time 170 s per tube). All sequencing was performed using an Illumina NextSeq 2000 machine.

### Computational analysis of methylated fractions

We developed two packages for methylated fraction analysis: snakemakeMethylFrac and methylFracAnalyzer. SnakemakeMethylFrac, a snakemake workflow ^44^, processes raw Illumina paired-end reads to determine methylated fractions at all GATC sites. Bowtie2 v2.5.1 ^45^ is used for alignment and bedtools v2.31.1 ^46^ is used to calculate the occupancy (fragment coverage) and 5’-end counting. GATC sites that overlap with CpG sites were filtered out, because DpnI cannot cut GATm^5^C. GATC half-sites that are within 150 bp of each other were also filtered out because small DNA fragments < 150 bp tend to be lost during sample purification. The output includes SQLite database and bigwig files. We use pandas ^47^, pyBigWig (https://github.com/deeptools/pyBigWig), biopython ^48^, matplotlib ^49^ and seaborn ^50^ in this workflow. MethylFracAnalyzer processes bigwig files from SnakemakeMethylFrac for downstream analysis. It calculates percentiles for each feature, methylation rates from median methylated fractions and relative methylation rates. It computes the average methylated fraction in 100-kb windows (T2T v1.1 assembly) and methylation rates using average methylated fractions. It calculates the average methylated fraction relative to the TSS of active and inactive genes, and relative to CTCF sites, smoothed in 21-bp windows. Finally, it generates the associated figures. This software uses pandas ^47^, pyBigWig, matplotlib ^49^, seaborn ^50^ and statsmodels ^51^.

### Analysis of RNA-seq, ATAC-seq and MNase-seq data

We used RNA-seq datasets from the GEO database (GSE201262 for MCF7 ^32^; GSE237066 for MCF10A ^52^) using salmon v1.10.0 ^53^ to normalise read counts. We averaged counts across replicates for each transcript using the GENCODE v43 annotation of the Hg38 assembly ^54^. For multi-transcript genes, we selected the most highly expressed transcripts and used their TSSs. We used published bigwig files for ATAC-seq datasets for MCF7 (GSE201262 ^32^) and MCF10A (GSE152410 ^55^). ATAC-seq data for 1-kb regions flanking TSSs were extracted; active genes were assigned based on average signal (>0.4 for MCF7 and >2 for MCF10A). Our MNase-seq data were aligned using Bowtie2 v2.5.1 ^45^. We selected read fragments for single nucleosomes (fragment length: 120-180 bp) and counted the nucleosome dyads in 2010-bp regions flanking active gene TSSs. We normalised dyad counts per gene using average dyad counts per flanking region, then averaged dyad counts across all active genes, smoothing in 21-bp windows.

### CTCF sites and ChromHMM

CTCF narrowPeaks were obtained from ENCODE ^56,57^ (ENCSR000AHD for MCF7 and ENCSR193SZD for MCF10A). CTCF motifs were predicted using HOMER v4.11.1 ^58^. For CTCF phasing analysis, the peaks containing a single copy of the highest frequency motif were selected. For ChromHMM analysis, we used ChIP-seq data from GSE85158 for both MCF7 and MCF10A cells ^38^, processed with the ENCODE ChIP-seq pipeline v2 (https://github.com/ENCODE-DCC/chip-seq-pipeline2). ChromHMM v1.23 predicted 15 chromatin states using the T2T v1.1 assembly ^39^. JHU RefSeqv110 + Liftoff v5.1 annotation (https://github.com/marbl/CHM13) were used for feature enrichment. Final chromatin state annotations are available in our GitHub repository (https://github.com/zhuweix/methylFracAnalyzer).

### Statistics and Reproducibility

Two biological replicate experiments were performed. The panels shown in each main figure belong to the same experiment. Extended Data figures generally show the results from both replicate experiments, except for immunoblots, micrographs and DNA gels. The immunoblots and DNA gel analyses were similar in both experiments. Correlations between biological replicates at the chromosomal level are presented in Extended Data Fig. 8.

## Acknowledgements

We thank Alex Vassilev for help with the FACS analysis. This study utilised the high-performance computational capabilities of the Biowulf Linux cluster at the National Institutes of Health (NIH). This research was supported by the Intramural Research Program of the NIH (NICHD).

## Author contributions

H.P. and P.E. performed the experiments; Z.X. performed the bioinformatic analysis; H.P. and D.C. wrote the manuscript.

## Competing interests

The authors have no competing interests.

Correspondence and requests for materials should be addressed to DJC.

---

**Extended Data Fig. 1.**
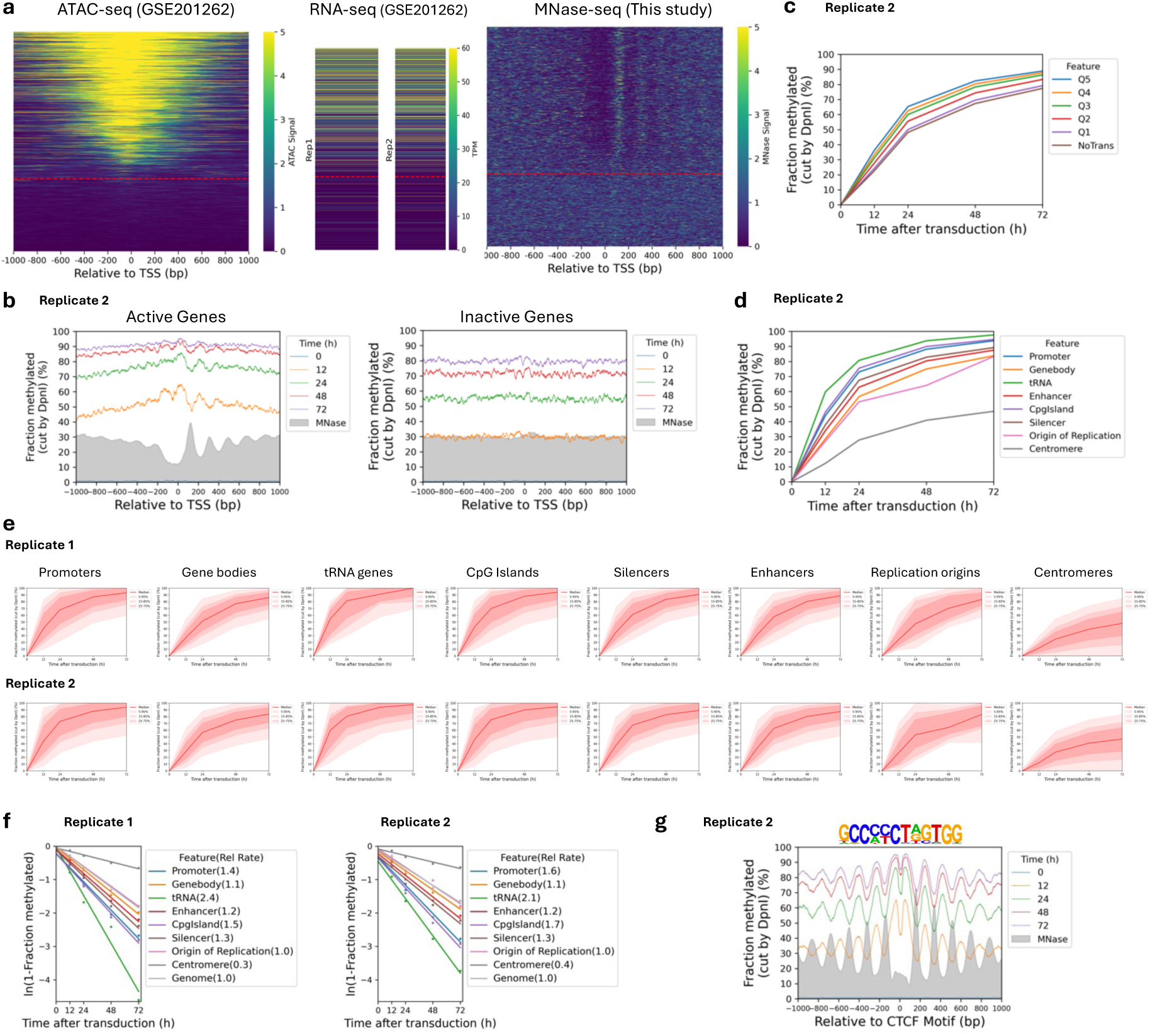
Rates of Dam methylation in various genomic regions in MCF7 cells. **a**, Left panel: All human genes sorted by ATAC-seq signal at their promoters in MCF7 cells ^32^ relative to the major TSS. The red line separates genes with NDRs (active) from those that have no NDR (inactive). Middle panel: RNA-seq data for MCF7 cells (from ^32^) sorted as in the left panel. Right panel: MNase-seq data sorted as in the left panel. **b,** Nucleosome phasing in vivo detected by Dam methylation for Replicate 2 (see Fig.1e,f for Replicate 1). Methylation data for GATC sites across active and inactive genes at each time point are plotted relative to the TSS (smoothed with a 21-bp window). Grey profile: nucleosome dyad distribution in nuclei (MNase-seq data normalised to 30%). **c,** Effect of transcription on median GATC site methylation. The active genes were divided into quintiles, Q1 to Q5, with increasing transcriptional activity; inactive genes were treated as a single group (“NoTrans”). Data for Replicate 2 (see Fig.1g for Replicate 1). **d,** Methylation time courses for the median GATC site in various genomic regions. Data for Replicate 2 (see Fig.1h for Replicate 1). **e,** Methylation time courses for various genomic regions defined by hg38 annotations. Red line: median GATC site; shading: data range as indicated. **f,** Relative methylation rates for various genomic regions in vivo. Rates are relative to the genomic average for all GATC sites. **g,** Nucleosome phasing around CTCF motifs in vivo using the motif shown. Data for Replicate 2 (see Fig.1i for Replicate 1).

**Extended Data Fig. 2.**
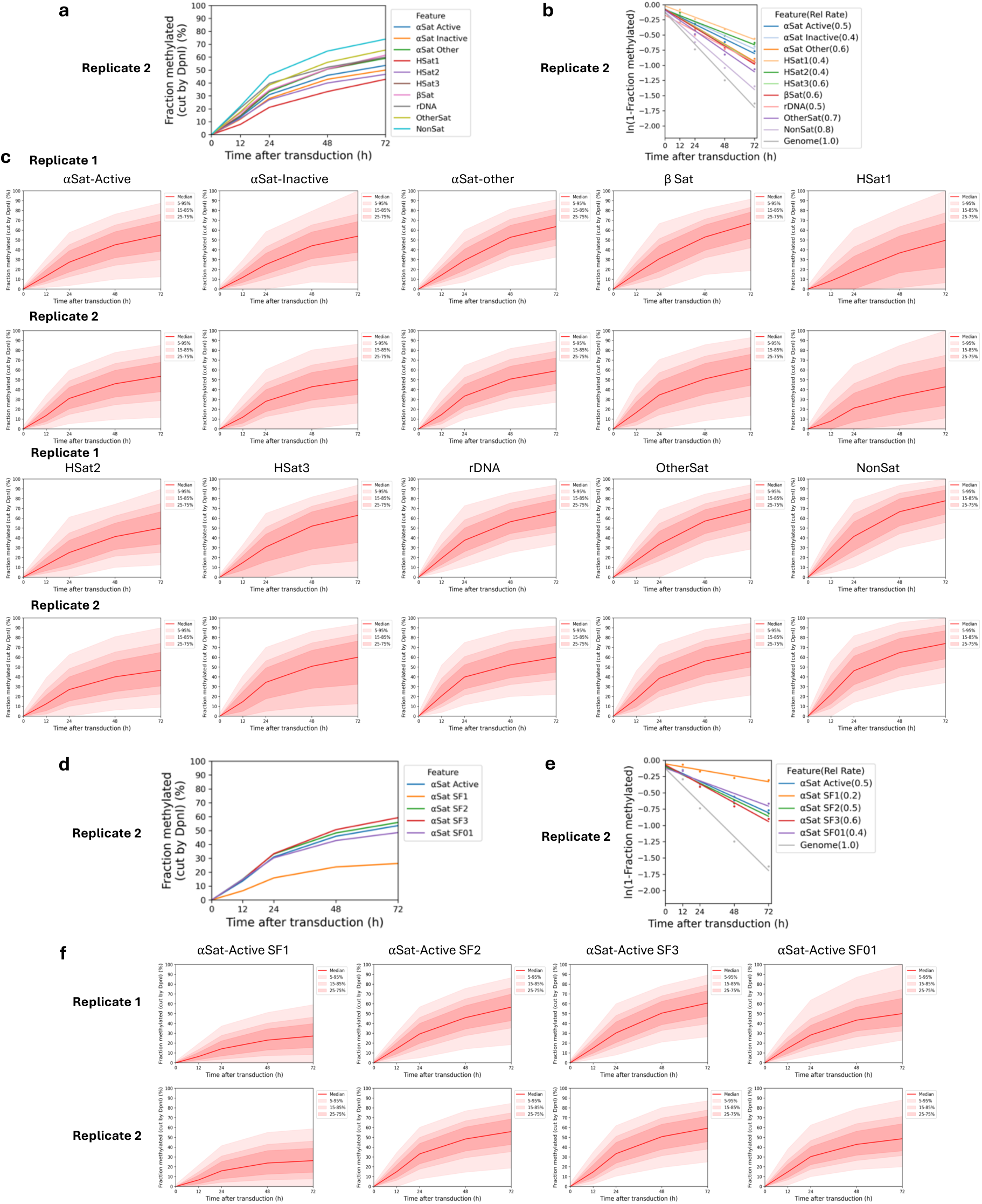
Dam methylation of the various centromeric satellite repeats in MCF7 cells. **a**, Methylation time courses for the median GATC site and **b,** methylation rates for the various centromeric elements relative to the genomic average site. Data for Replicate 2 (see Fig. 2c,d for Replicate 1). **c,** Methylation time courses for the median GATC site for the various centromeric elements. Red line: median GATC site; shading: data range as indicated. **d,** Methylation time courses for the median GATC site and **e,** relative methylation rates for the various active α-satellite supra-chromosomal families. Data for Replicate 2 (see Fig. 2e,f for Replicate 1). **f,** Methylation time courses for the median GATC site for the various active α-satellite supra-chromosomal families. Red line: median GATC site; shading: data range as indicated.

**Extended Data Fig. 3.**
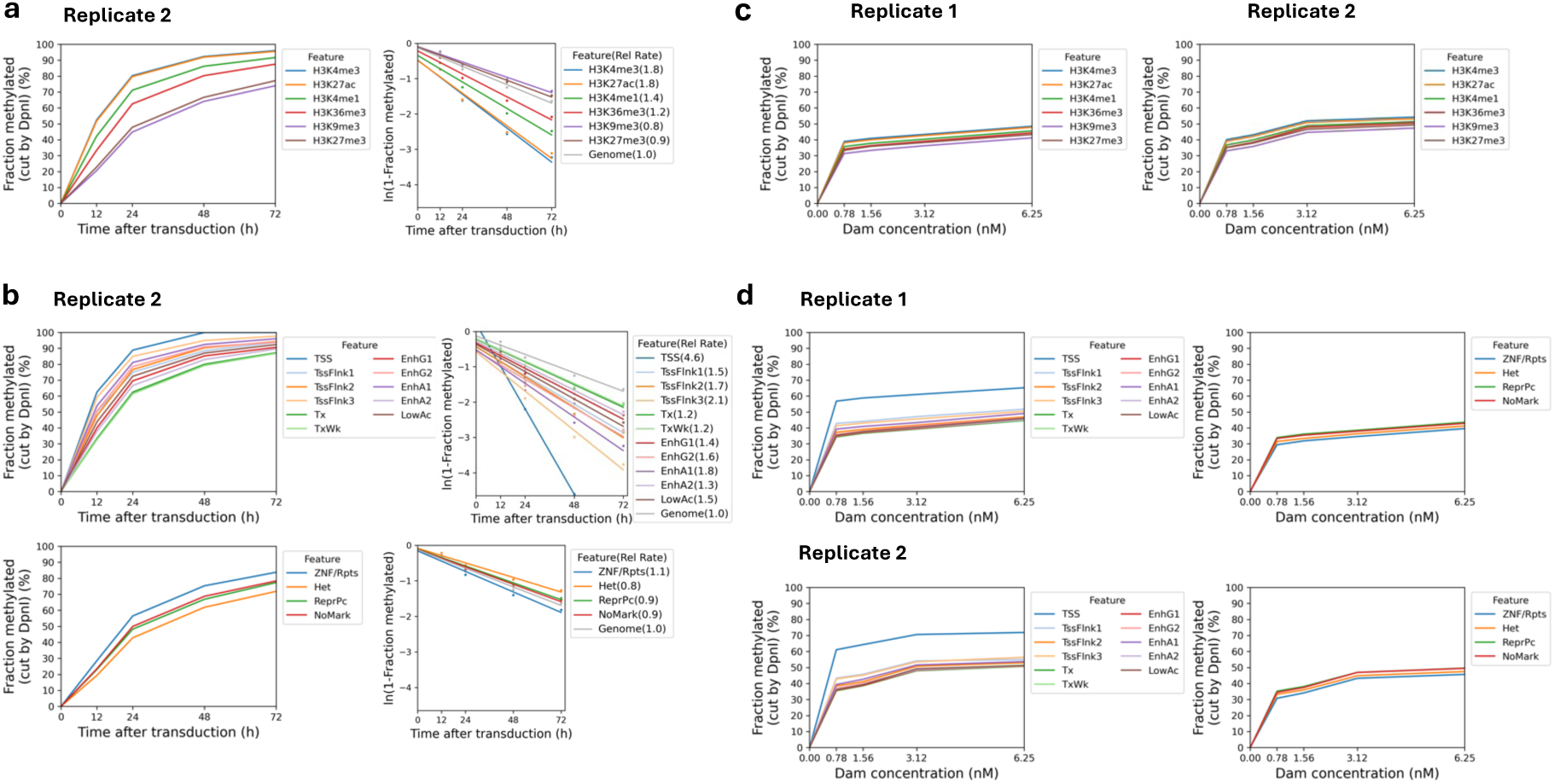
Dam methylation of heterochromatin and euchromatin in living MCF7 cells and in MCF7 nuclei. **a**, Methylation time courses for the median GATC site and methylation rates for regions marked by histone modifications associated with euchromatin (H3K4me1, H3K4me3, H3K27ac or H3K36me3) or heterochromatin (H3K9me3 or H3K27me3), relative to the genomic average site. ChIP-seq data from ^38^. Data for Replicate 2 (see Fig. 3a,b for Replicate 1). **b,** Methylation time courses for the median GATC site and relative methylation rates for the 15 epigenetic states defined by our ChromHMM model (see Fig. 3b). Data for Replicate 2 (see Fig. 3c,d for Replicate 1). **c,** Methylation of the median GATC site in nuclei for regions marked by histone modifications associated with euchromatin or heterochromatin. **d,** Dam methylation in isolated nuclei of the different chromatin states specified by the ChromHMM model (see Fig. 3b).

**Extended Data Fig. 4.**
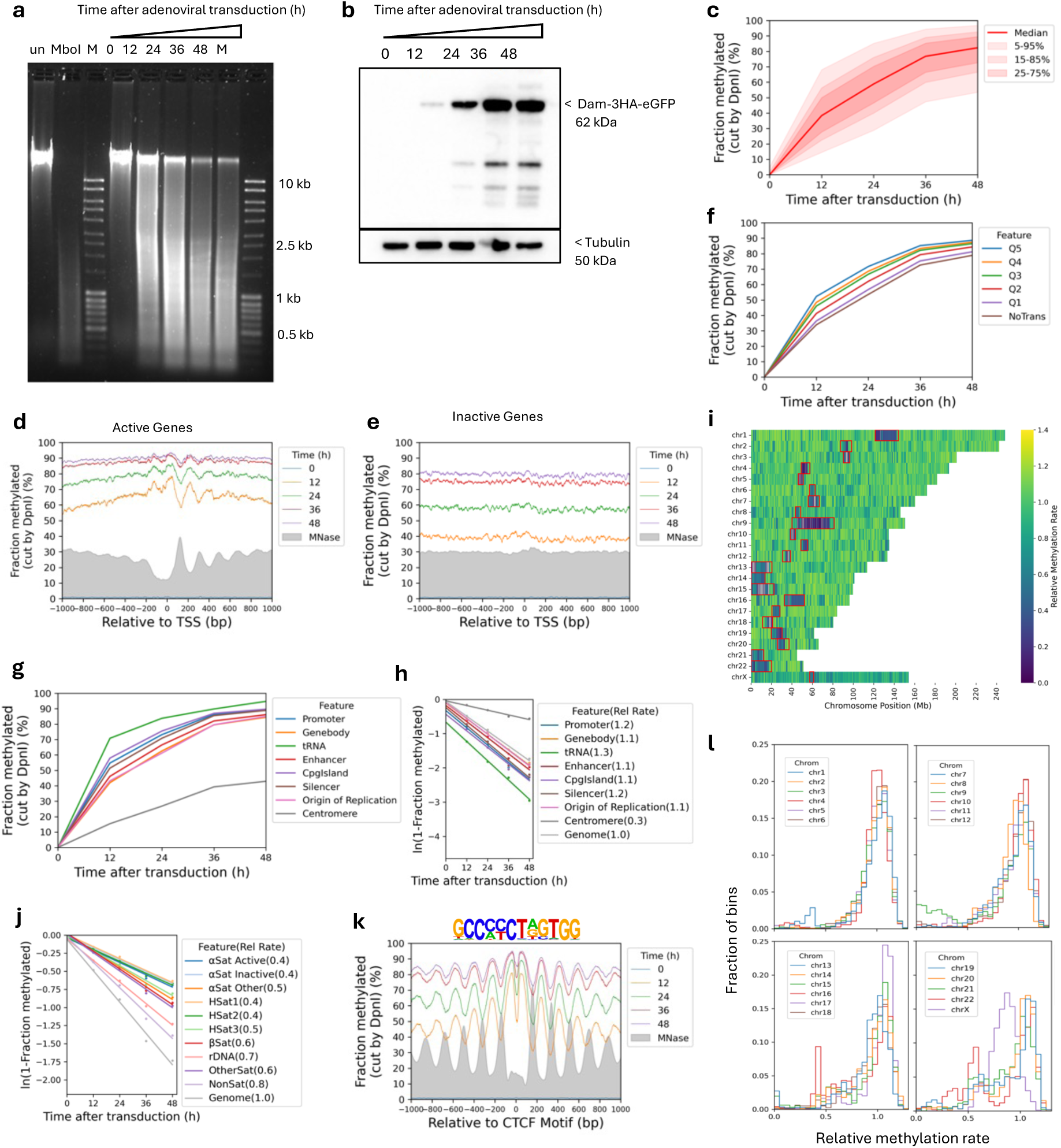
The human genome is globally accessible in live MCF10A cells. **a**, Agarose gel electrophoresis of DpnI-digested genomic DNA purified from MCF10A cells as a function of time of adenovirus treatment. ‘un’, undigested genomic DNA; ‘MboI’, DNA from non-transduced cells digested with MboI; M, DNA size marker. **b,** Anti-HA immunoblot to detect Dam-3HA-eGFP expression in MCF10A cells. **c,** Almost complete methylation of GATC sites in MCF10A cells after transduction. Red line and shading: median GATC site methylation with data range indicated. **d,e,** Nucleosome phasing with respect to the TSS for active and inactive genes, as defined by ATAC-seq data for MCF10A cells 55. Grey profile: nucleosome dyad distribution in nuclei (MNase-seq data for MCF7 cells arbitrarily normalised to 30%). **f,** The effect of transcriptional activity on methylation rate. Active genes were divided into quintiles Q1 to Q5 based on increasing transcriptional activity (Q5 is the highest) using RNA-seq data for MCF10A cells from (Dorgham et al. 2023); methylation of the median GATC site in each quintile is shown. Inactive genes are treated as a single separate group (’NoTrans’). **g,** Median GATC methylation for various genomic regions using annotations from the hg38 genome. **h,** Relative median GATC site methylation rates for various genomic regions. **i,** Heat map showing the variation in methylation rate at the chromosomal level in MCF10A cells. The average methylation rate was calculated for all GATC sites in each 100 kb window in the T2T genome by plotting ‘ln (1-fraction methylated)’ against time after adenovirus transduction, and then normalised to the genomic average rate to obtain relative rates. Red rectangles: centromeric regions. **j,** Relative methylation rates for the various centromeric elements. **k,** Nucleosome phasing around CTCF motifs in MCF10A cells using the motif shown. **l,** Relative methylation rate data derived as in ‘i’ for each of the 23 chromosomes are separated into four separate plots for ease of comparison. Histograms of the fraction of 100-kb windows having a given relative methylation rate.

**Extended Data Fig. 5.**
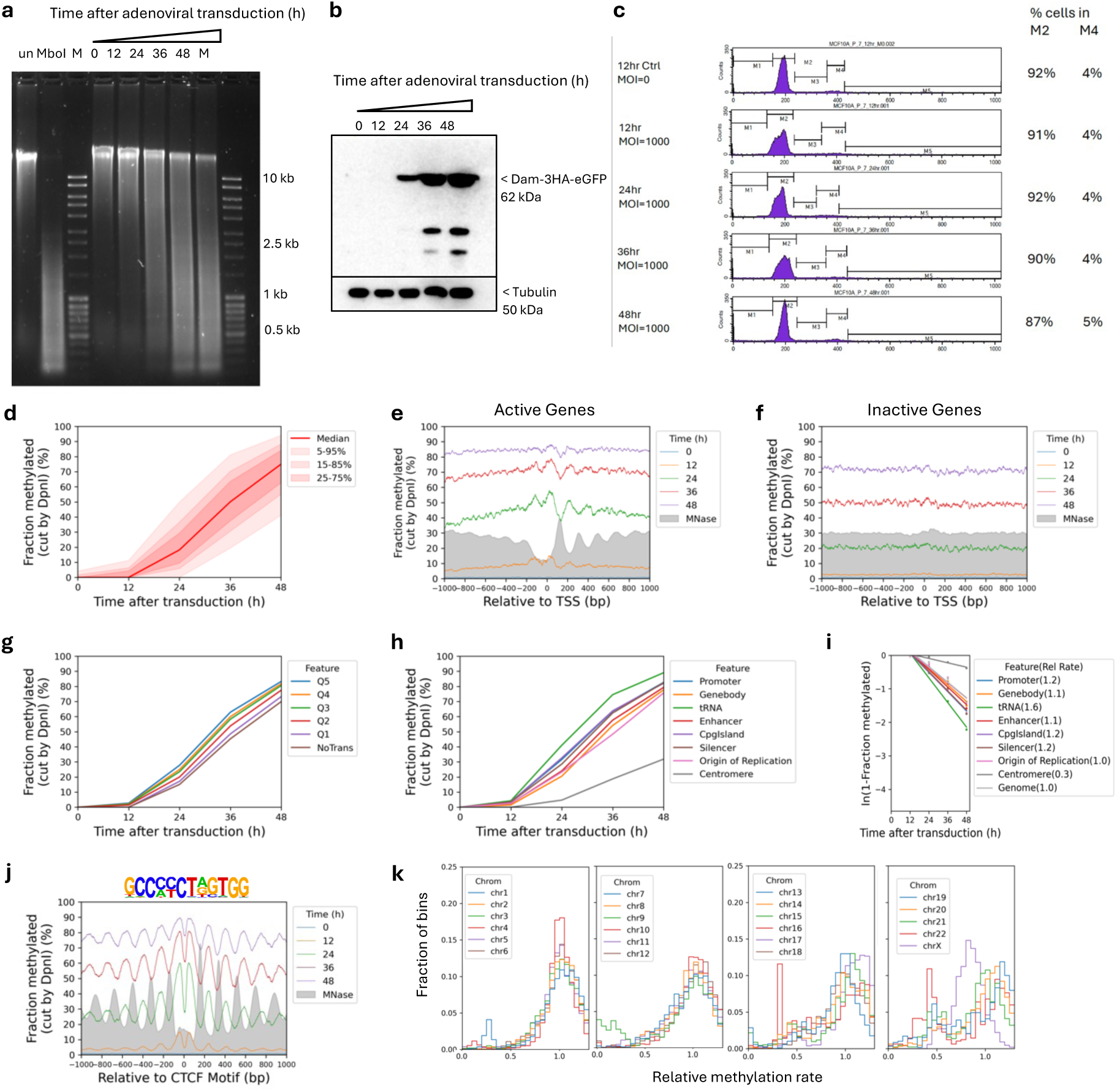
The human genome is globally accessible in confluent MCF10A cells. **a**, Agarose gel electrophoresis of DpnI-digested genomic DNA purified from confluent MCF10A cells as a function of time of adenovirus treatment. ‘un’, undigested genomic DNA; ‘MboI’, DNA from non-transduced cells digested with MboI; M, DNA size marker. **b,** Anti-HA immunoblot to detect Dam-3HA-eGFP expression in MCF10A cells. **c,** FACS analysis confirms that the confluent MCF10A cells are arrested in G1 during the time course after adenovirus transduction. **d,** Time course of Dam methylation of all GATC sites in MCF10A cells after transduction. Red line and shading: median GATC site methylation with data range indicated. **e,f,** Nucleosome phasing with respect to the TSS for active and inactive genes, as defined by ATAC-seq data for MCF10A cells ^55^. Grey profile: nucleosome dyad distribution in nuclei (MNase-seq data for MCF7 cells arbitrarily normalised to 30%). **g,** The effect of transcriptional activity on methylation rate. Active genes were divided into quintiles Q1 to Q5 based on increasing transcriptional activity (Q5 is the highest) using RNA-seq data for MCF10A cells (Dorgham et al. 2023); methylation of the median GATC site in each quintile is shown. Inactive genes are treated as a single separate group (’NoTrans’). **h,** Median GATC methylation for various genomic regions using hg38 genome annotations. **i,** Relative median GATC site methylation rates for various genomic regions. **j,** Nucleosome phasing around CTCF motifs in MCF10A cells using the motif shown. **k,** Histograms of the fraction of 100-kb windows having a given relative methylation rate for each of the 23 chromosomes are separated into four separate plots for ease of comparison.

**Extended Data Fig. 6.**
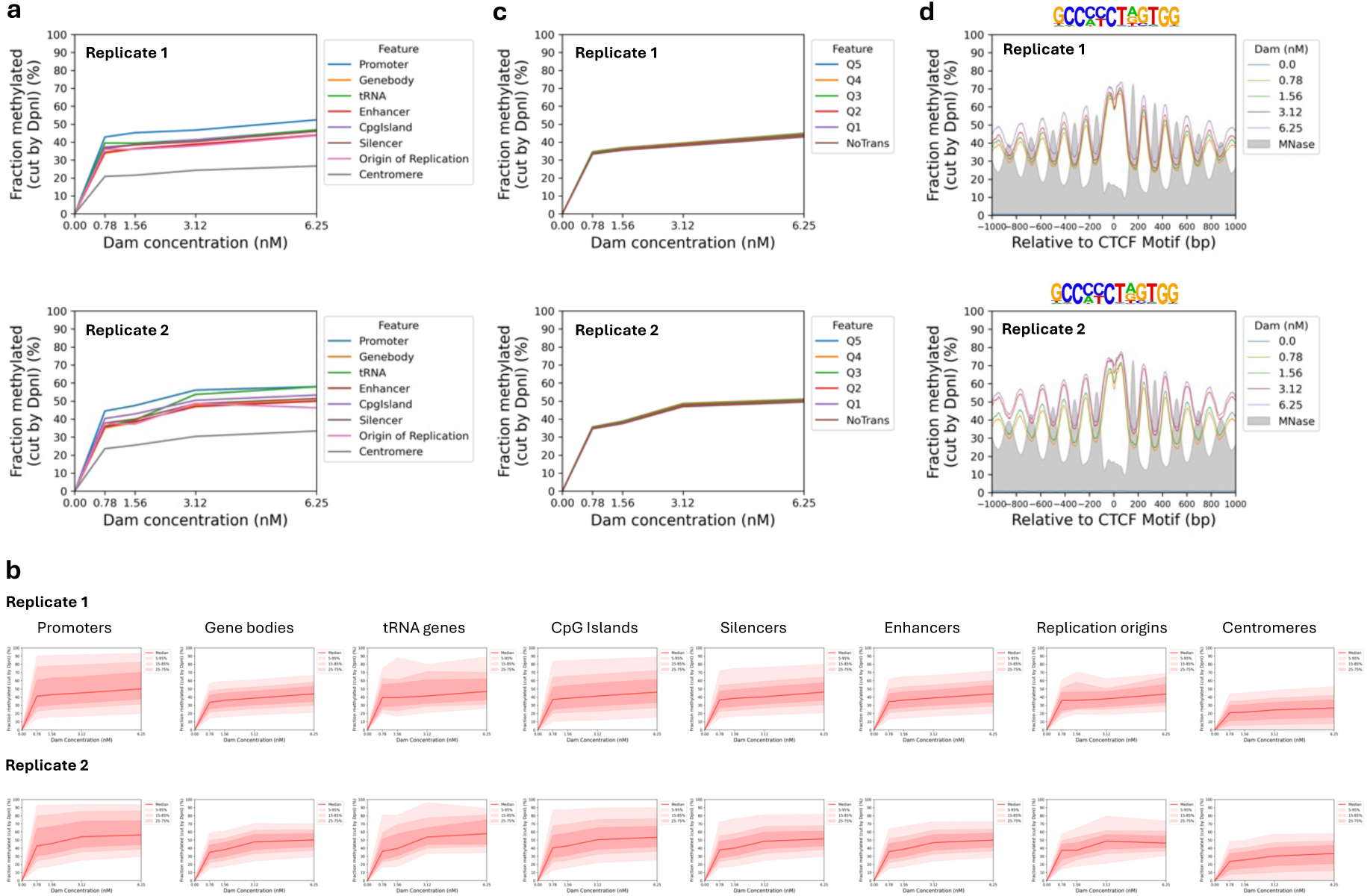
Limited genome accessibility in isolated MCF7 nuclei. Nuclei were treated with increasing concentrations of Dam. **a,** Comparison of the methylation of the median GATC site in various genomic regions as a function of Dam concentration. **b,** Separate plots for methylation of the median GATC site in various genomic regions as a function of Dam concentration. Red line and shading: median GATC site methylation with data range indicated. **c,** The effect of transcriptional activity on methylation rate in nuclei. Active genes were divided into quintiles Q1 to Q5 based on increasing transcriptional activity (Q5 is the highest) using RNA-seq data for MCF7 cells ^32^; methylation of the median GATC site in each quintile is shown. Inactive genes are treated as a single separate group (’NoTrans’). **d,** Nucleosome phasing in nuclei relative to CTCF motifs detected by Dam methylation using the motif shown. Methylation data for GATC sites at each Dam concentration are plotted relative to each CTCF site (smoothed with a 21-bp window). Grey profile: nucleosome dyad distribution in nuclei (MNase-seq data normalised to 30%).

**Extended Data Fig. 7.**
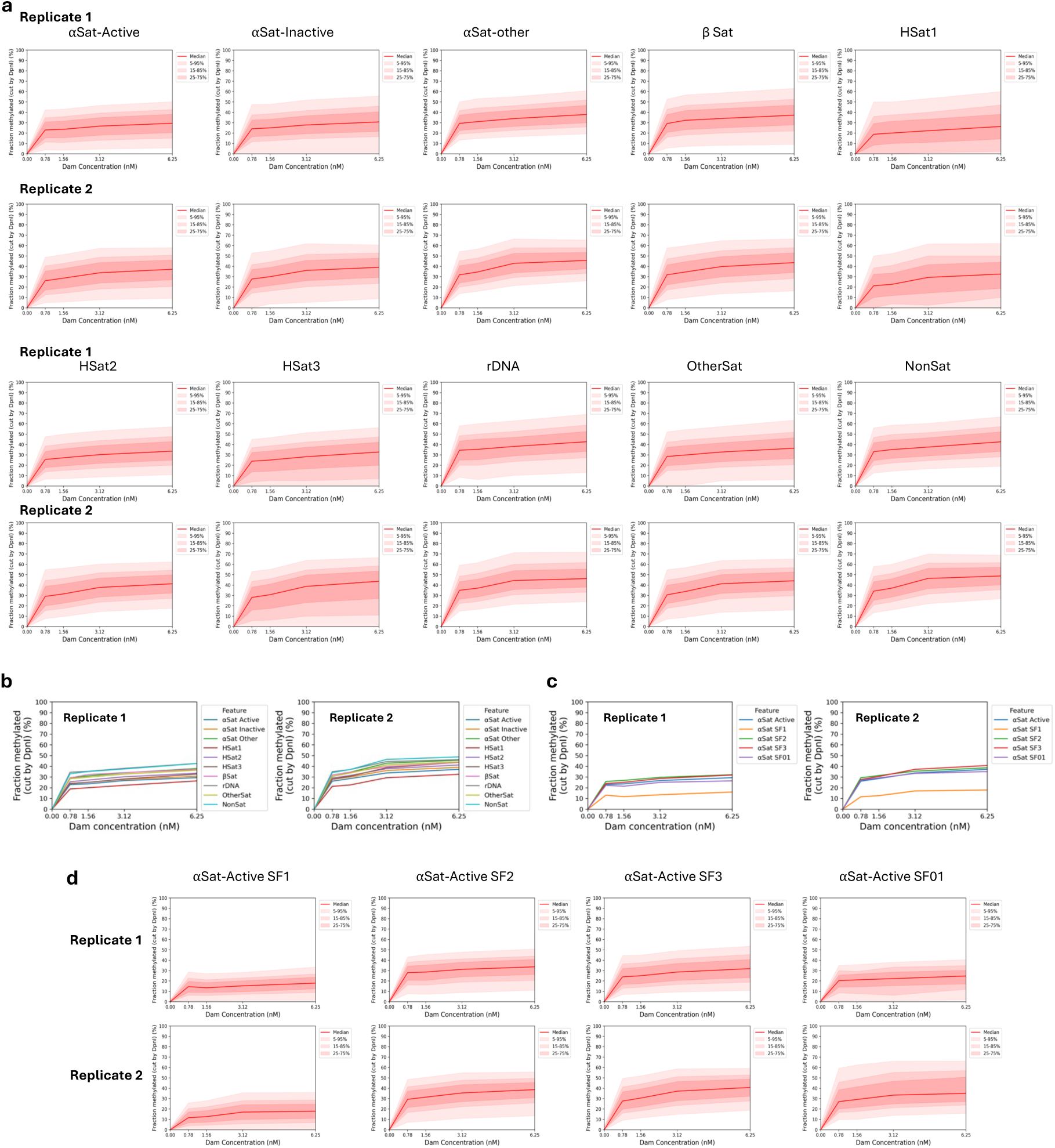
Methylation of centromeric elements is limited in nuclei. **a**, Separate plots showing methylation of the median GATC site in various centromeric elements as a function of Dam concentration. Red line and shading: median GATC site methylation with data range indicated. **b,** Comparison of the methylation of the median GATC site in the various centromeric elements as a function of Dam concentration. **c,** Comparison of the methylation of the median GATC site in the various active α-satellite supra-chromosomal families as a function of Dam concentration. **d,** Separate plots showing methylation of the median GATC site in various active α-satellite supra-chromosomal families as a function of Dam concentration.

**Extended Data Fig. 8.**
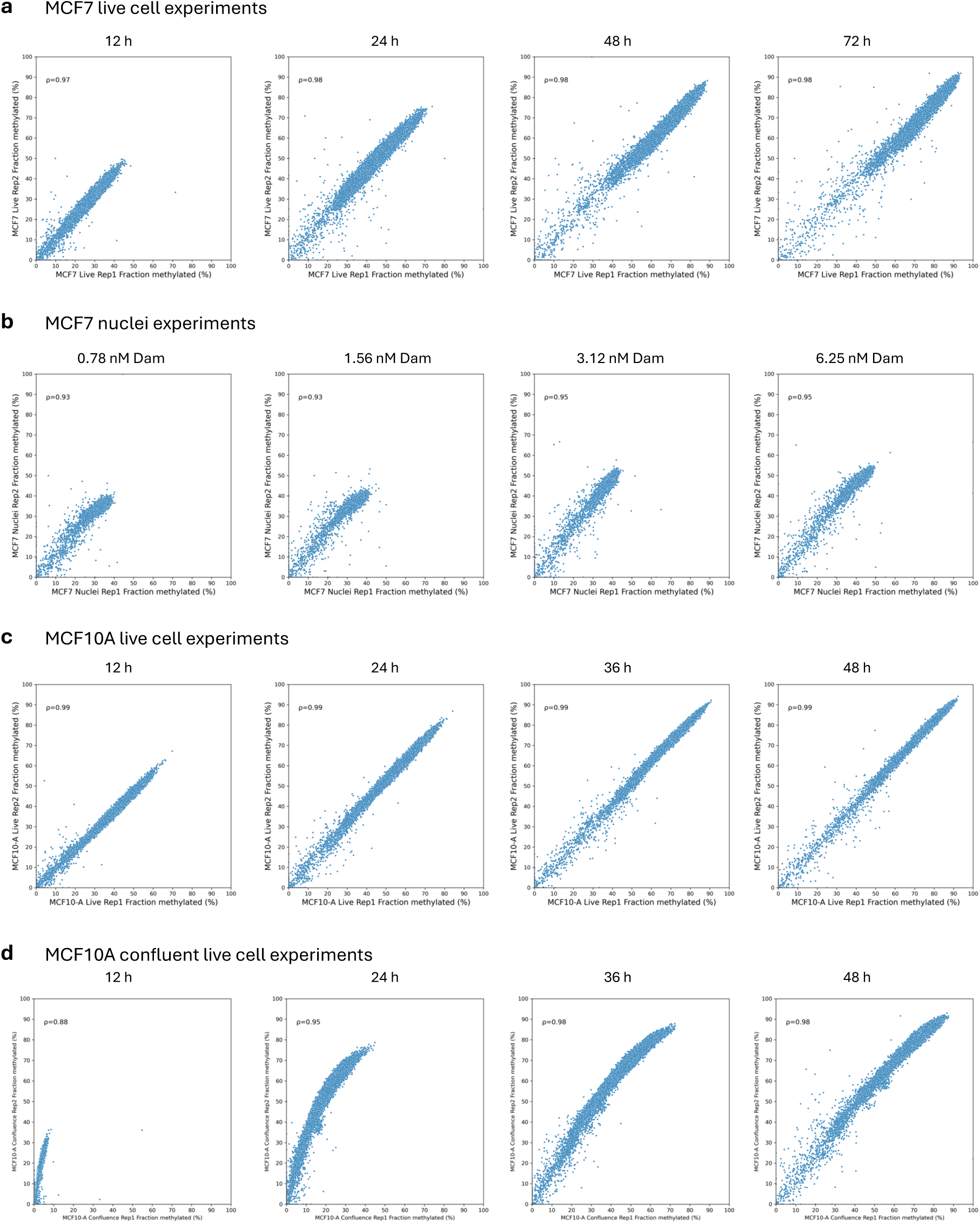
Comparison of biological replicate experiments at the chromosomal level. Pearson correlations for the average % methylated for all GATC sites in each 100 kb window for each time point (live cells) or Dam concentration (nuclei). **a,** MCF7 cells, **b,** MCF7 nuclei, **c,** Dividing MCF10A cells, **d,** Confluent MCF10A cells.

